# Pattern formation and travelling waves in a multiphase moving boundary model of tumour growth

**DOI:** 10.1101/2022.11.23.517688

**Authors:** Jacob M. Jepson, Reuben D. O’Dea, John Billingham, Nabil T. Fadai

**Affiliations:** School of Mathematical Sciences, University of Nottingham, Nottingham, NG7 2RD, UK

**Keywords:** Stability analysis, Asymptotic analysis, Moving boundary problem, Tissue mechanics

## Abstract

We analyse a multiphase, moving boundary model that describes solid tumour growth. We consider the evolution of a motile, viscous cell phase and an inviscid extracellular liquid phase. The model comprises two partial differential equations that govern the cell volume fraction and the cell velocity, together with a moving boundary condition for the tumour edge. Numerical simulations of the model indicate that patterned solutions can be obtained, which correspond to multiple regions of high cell density separated by regions of low cell density. In other parameter regimes, solutions of the model can develop into a forward- or backward-moving travelling wave, corresponding to tumour growth or extinction, respectively. A travelling-wave analysis allows us to find the corresponding wave speed, as well as criteria for the growth or extinction of the tumour. Furthermore, a stability analysis of these travelling-wave solutions provides us with criteria for the occurrence of patterned solutions. Finally, we discuss how the initial cell distribution, as well as parameters related to cellular motion and cell-liquid drag, control the qualitative features of patterned solutions.

## 1 Introduction

A tumour is characterised by an abnormal mass of tissue that develops when cells divide more frequently or die less frequently than they should. Benign tumours are non-invasive and remain local to the site at which they first developed, whilst malignant (cancerous) tumours possess uncontrollably dividing cells that can spread through the body (Patel, 2020). In the initial stages of growth, the tumour is sufficiently small that cells are adequately nourished by nutrient obtained from the surrounding tissue by diffusion. As the tumour grows, a limited amount of nutrient enters the core due to its consumption by cells in the tumour rim, and malnourished cells can release agents that stimulate angiogensis (Fam et al., 2003). Consequently, the new blood vessels formed by angiogensis provide a means for metastasis, whereby tumour cells enter the blood stream and spread through the body to form secondary tumours (Zetter, 1998). The effect that cancer has on the global population encourages vigorous scientific research into its treatment and prevention.

Extensive mathematical research has been undertaken to better understand tumour growth. Some of the earliest mathematical models of tumour growth examine the diffusive nature of substances moving through cells in order to investigate how physiological processes such as oxygen transport and waste disposal affect tumour growth (Hill, 1928; Thomlinson and Gray, 1955; Burton, 1966). For example, the study Greenspan (1972) captures the principal structure of an avascular tumour by incorporating the effects of a diffusive nutrient and a growth inhibitor. More recent studies analyse how cells migrate within a tumour in response to various stimuli (McElwain and Pettet, 1993; Thompson and Byrne, 1999). For example, Pettet et al. (2001) determines how the concentration of nutrient present at the tumour edge affects the migration of cancer cells in different regions of the tumour. In contrast to these earlier models of tumour growth, Ward and King (1997) derives a system of partial differential equations to track the behaviour of living and dead cells, whereby the volume exchange between the two cell species drives the velocity field within the tumour. Some authors adopt a microscale approach, which seeks to model interactions between a large number of individual cells. Whilst such approaches can formalise single-cell dynamics observed experimentally, they can can become computationally infeasible for tissue-scale simulations (Poleszczuk and Enderling, 2014). Reviews of microscale approaches to model tumour growth via cellular automaton and agent-based systems is provided in Boonderick et al. (2010) and Macnamara (2021), respectively.

Conversely, and of greater relevance to this study, some authors adopt a multiphase approach to model the growth of a solid tumour or a tissue construct (Jepson et al., 2022; O’Dea et al., 2010; Tosin and Preziosi, 2010; Sciumè et al., 2013). In contrast to continuum approaches which model tissue as a homogeneous mass, such as Greenspan (1972), multiphase models allow the interaction of different tissue constituents, and have been proposed to be a more natural modelling framework for studying solid tumour growth than existing theories. For example, Byrne et al. (2003) develops a multiphase model which describes the interaction between a cell and a liquid phase in an avascular tumour; a detailed analysis of this model is provided in Breward et al. (2002), and the cell phase in this model exhibits either travelling waves which propagate with constant speed or a steady state, both of which are in agreement with experimental observations. In Lemon and King (2007), a multiphase, moving boundary model of tissue growth is analysed to determine how mechanical pressures influence the attractive or dispersive nature of cells. In certain parameter regimes, the tissue evolves at a constant speed as either a forward or backward travelling wave. Alternatively, solutions can exhibit patterned solutions, whereby cells form distinct regions of high-cell density that are separated by regions of low cell density. Another multiphase description of tissue growth which admits patterned solutions is presented in Green et al. (2018), where cells can migrate toward a concentration of nutrient and form regions of high cell density. To identify parameter regimes for which pattern formation is expected, Green et al. (2018) employs a Turing-type stability analysis (Turing, 1952) and determines when a spatially-uniform steady state is unstable to small-amplitude perturbations.

In a similar fashion to Green et al. (2018), many other mathematical models employ well-established techniques to analyse the existence of patterned solutions on a fixed spatial domain; see Murray (2002) for a review. In contrast, mathematical models have been proposed to examine how patterned solutions are affected by an evolving spatial domain (Crampin et al., 2002; Krause et al., 2018; Toole and Hurdal, 2014). While moving boundary models can provide a more realistic description of some biological phenomena than fixed domain models, the stability analysis exploited to obtain criteria for patterned solutions is generally more complex due to the dependence of any spatially homogeneous base states on time (Gorder et al., 2021). To overcome this difficulty, Gorder et al. (2021) uses a comparison principle to provide criteria for spatial patterning in terms of differential inequalities, for a class of spatially evolving reaction-diffusion systems.

In this work, we analyse patterned and travelling-wave solutions of the multiphase, moving boundary model developed in Byrne et al. (2003). This model describes solid tumour growth, and considers the evolution of a motile, viscous cell phase and an inviscid extra-cellular liquid phase, both of which are modelled as incompressible fluids and nutrient-rich. Tissue mechanics, cellular growth and a mechanism to represent cell-liquid drag are accounted for by considering relevant constitutive assumptions in a similar fashion to those in Byrne et al. (2003) and Breward et al. (2002). Following King and Franks (2004), we assume that nutrient is abundantly distributed throughout the tumour. In the context of *in vivo* tumour growth, this assumption is physically relevant where the tumour is in the initial stage of growth and all cells are adequately nourished (Franks and King, 2003). This nutrient-rich assumption is also appropriate when considering the initial growth of a suspension of *in vitro* tumour cells (Byrne et al., 2003). Whilst the model developed in Byrne et al. (2003) pertains to both *in vivo* and *in vitro* tumour growth, we emphasise that the mathematical results obtained in its analysis can be applied to a wider class of multiphase tissue growth models, such as that in Lemon and King (2007).

The paper is constructed as follows. In Section 2, the model from Byrne et al. (2003) is summarised and dimensionless equations of mass transfer and momentum balance are stated. Following this, some exemplar numerical solutions of the model are presented in Section 3, which exhibit patterned solutions and forward- and backward-moving travelling waves. In Section 4, a travelling-wave analysis is presented. This allows us to find the speed of travelling waves, as well as criteria for the growth or extinction of the tumour. We also present a stability analysis of these travelling-wave solutions and obtain criteria for when patterned solutions can occur. In Section 5, we neglect the effect of the moving tumour edge and determine the stability of a spatially-uniform steady state. A comparison between the results of the travelling-wave and spatially-uniform stability analysis allows us to suggest that the moving boundary does not contribute toward the formation of spatial patterns. In Section 6, we examine the qualitative features of patterned solutions. We find that the initial cell distribution and the value of the cell-liquid drag have a significant effect on the features of patterns, in comparison to the strength of forces generated by cellular motion. In Section 7, we discuss the behaviour of the model and highlight the mathematical and biological results obtained in its analysis.

## 2 Model development

In this section, we present a summary of the two-phase model developed in Byrne et al. (2003) which describes the growth of a solid tumour. Following King and Franks (2004), we assume that nutrient is abundantly distributed throughout the tumour; see Section 1 for a discussion regarding the biological relevance of this assumption. For simplicity, we formulate the model in a one-dimensional Cartesian geometry and present the model in dimensionless form for brevity.

The tumour model of Byrne et al. (2003) consists of two phases, denoted by *n*(*x, t*) and *w*(*x, t*), that represent the volume fraction of cells and extracellular liquid, respectively. These phases satisfy the no-voids volume constraint *n* + *w* = 1. The velocity fields *v*_*n*_(*x, t*) and *v*_*w*_(*x, t*), as well as the pressures *p*_*n*_(*x, t*) and *p*_*w*_(*x, t*), are associated with the phases *n* and *w*, accordingly. We model the cell and liquid phases as viscous and inviscid fluids, respectively. The spatial domain of the tumour evolves over time due to cellular motion, so the volume fractions *n* and *w* evolve on the moving domain 0 ≤ *x* ≤ *L*(*t*), where *x* = 0 and *x* = *L*(*t*) denote the tumour core and tumour edge, respectively. The model is developed by considering mass and momentum balances for each phase, assuming that the phases are incompressible with equal density, and by neglecting inertial effects.

The mass transfer equation for the cell phase is represented by the partial differential equation (PDE)

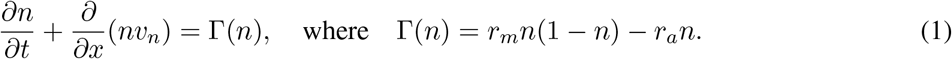

Here, *w* has been replaced with 1 − *n*, and *r*_*m*_ and *r*_*a*_ are positive constant dimensionless rates of cell mitosis and apoptosis, respectively. Equation (1) states that the rate of change of the cell volume fraction *n* is due to advection, as well as the production and death of cells. Since daughter cells are constructed via mitosis using the available liquid, the first term on the right hand side of Γ(*n*) is proportional to the liquid volume fraction.

The momentum balance equation for the cell phase is

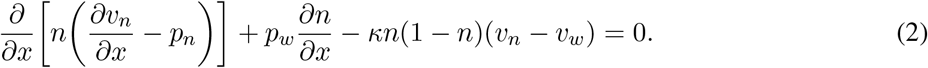

The first term in (2) describe stresses in the cell phase and consists of contributions from the cell viscosity and pressure in the cell phase. The third term in (2) describes the forces exerted on the cells by the pressure in the liquid. The fourth term represents the interphase drag between the cells and liquid, where *κ* is the dimensionless drag coefficient. Combining (2) with the momentum balance equation for the liquid phase and the overall conservation of mass condition allows the elimination of *v*_*w*_, *p*_*n*_ and *p*_*w*_, and provides

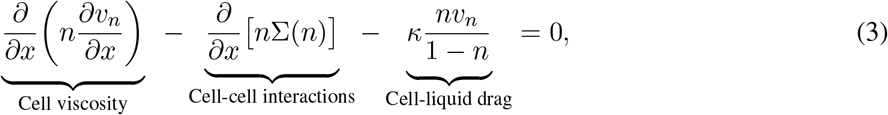

where Σ(*n*) = *p*_*n*_ − *p*_*w*_ represents the additional pressures that arise due to cell-cell interactions.

Following Byrne et al. (2003) and Green et al. (2009), an appropriate expression for Σ(*n*) is

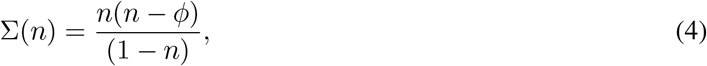

where 0 *< ϕ <* 1 is the natural packing density of the cells. When *n > ϕ*, the cells membranes experience stress, and the cells will repel each other to relieve it. This repulsion becomes large as the available space for the cells decreases, which is reflected in the singularity at *n* = 1. When *n < ϕ*, the cells will attract one another, due to their filopodia coming into contact (Breward et al., 2002). Necessarily, we have Σ(0) = 0.

Equations (1) and (3) are coupled to the boundary conditions

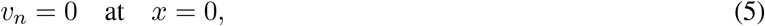

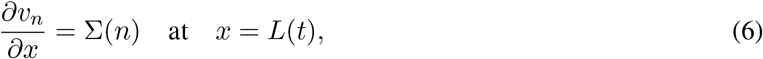

which arise from the assumptions that the tumour is symmetric about its centre (*x* = 0), and that the tumour edge is stress-free and *p*_*w*_(*L, t*) = 0. The moving boundary at *x* = *L*(*t*) moves with the cell velocity, so that

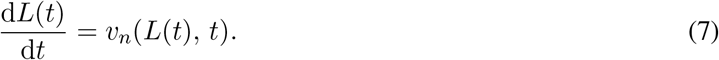

The initial distribution of *n* and position of the tumour boundary are

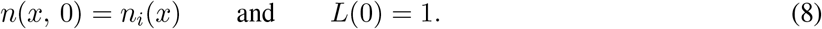

Various choices for *n*_*i*_(*x*) are made throughout the paper.

For clarity, the model consisting of equations (1), (3), (5)**–**(8) will be henceforth referred to as the moving boundary model (MBM). Throughout this paper, we assume that *r*_*a*_ *< r*_*m*_ so that there is net cell growth. As shown in Byrne et al. (2003) and Breward et al. (2002), combining (1), (6) and (7) at *x* = *L*(*t*) provides the autonomous ordinary differential equation (ODE)

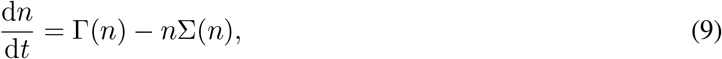

so that if *r*_*m*_ *> r*_*a*_, then *n*(*L, t*) ~ *n*_∞_ for sufficiently large time, where Γ(*n*_∞_) − *n*_∞_Σ(*n*_∞_) = 0. In subsequent large-time analysis, we may therefore replace the boundary condition from (6) with *n*(*L, t*) = *n*_∞_.

## 3 Numerical results

To illustrate the behaviour of the MBM, we present and discuss some numerical solutions on the moving domain 0 ≤ *x* ≤ *L*(*t*). In this section we fix *r*_*m*_ = 0.3, *r*_*a*_ = 0.2 and *κ* = 100, and pay particular attention to three exemplar values of *ϕ* that generate patterned and forward- and backward-moving travelling-wave solutions. Since the tumour is nutrient-rich and all cells are adequately nourished, we take *n* to be near-uniform when *t* = 0, and adopt the initial conditions 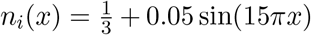. This also accounts for small fluctuations in the cell density across the tumour. We note that the choice of 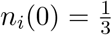 arises from the far-field value of *n* in travelling-wave coordinates as considered in Section 4, and is chosen here for convenience.

To obtain numerical solutions of the MBM, we fix the moving boundary by scaling *x* with *L*(*t*) as *ξ* = *x/L*(*t*), so that *ξ* ∈ [0, 1], and the model becomes

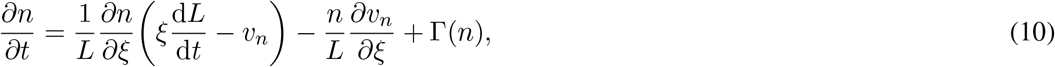

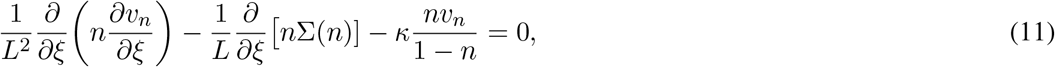

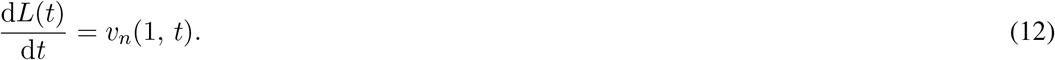

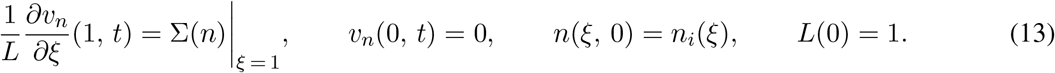

We spatially discretise (10) and (11) using first-order finite differences. Upwind or downwind finite differences are used for the first term on the right-hand side of (10), the direction of which is determined by the sign of the quantity in the brackets. The equations from (10) and (12) are then numerically integrated in time using the function `ode23s` in MATLAB, which uses a third order Runge-Kutta method.

In Fig. 1(a, b), we present *n* and *v*_*n*_ when *ϕ* = 0.2, where we observe forward-moving travelling waves and linear growth in *L*, after an initial period of transient growth from the initial cell distribution. In contrast, Fig. 1(c, d) illustrate *n* and *v*_*n*_ when *ϕ* = 0.5, where we observe backward-moving travelling waves and linear recession in *L*. As the tumour vanishes, the tumour edge decays exponentially towards zero, as shown in Fig. 1(d). Clearly, the case in which *ϕ* = 0.2 corresponds to tumour growth, whereas *ϕ* = 0.5 corresponds to the retreat and eventual extinction of the tumour. This motivates a travelling-wave analysis of the MBM which is provided in Section 4, where we express the speed of the tumour edge in terms of the model parameters and obtain explicit criteria for whether the wave grows or retreats.

**Figure 1:**
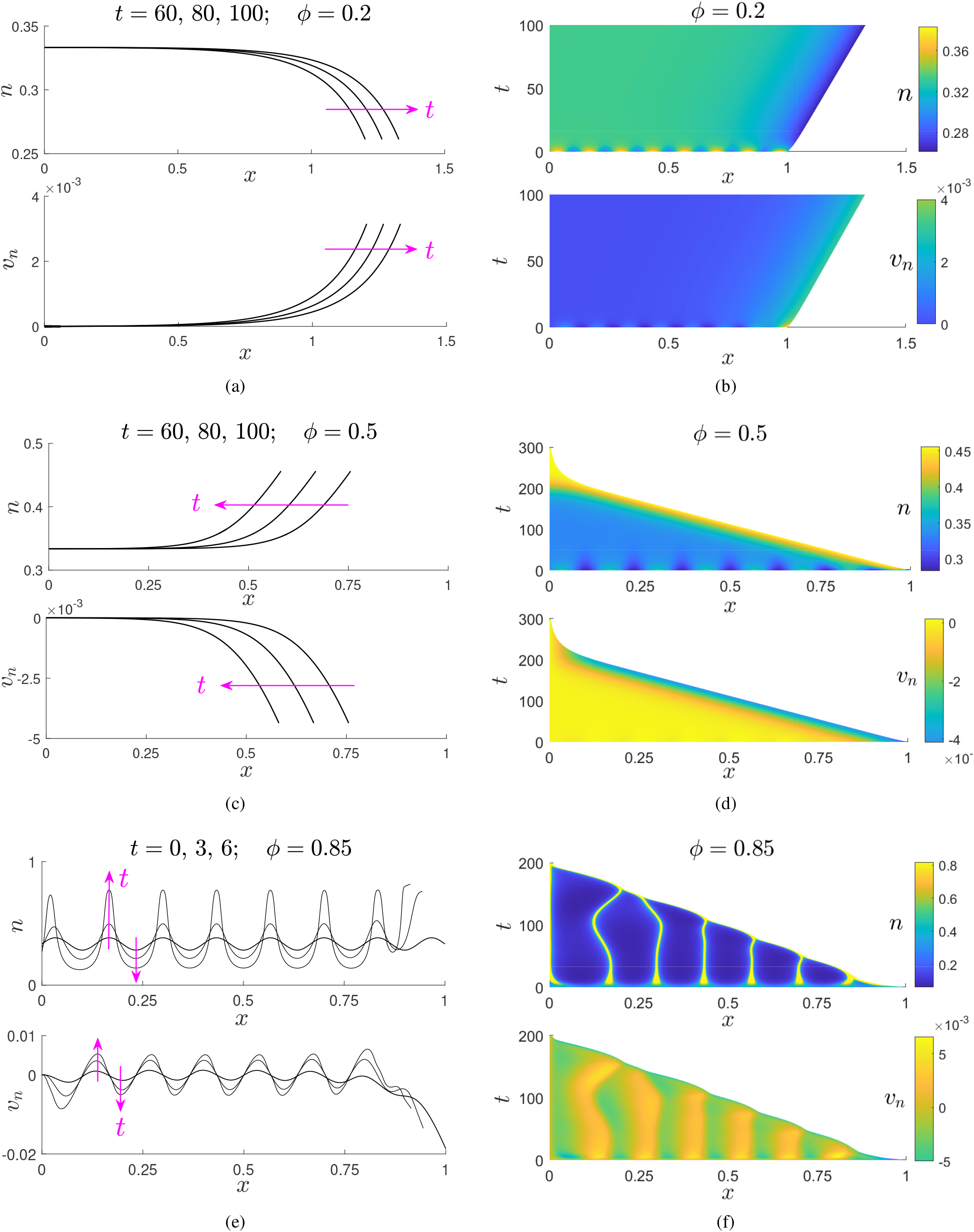
Numerical solutions of the MBM for three values of *ϕ*. The black lines in (a, c, e) represent *n*(*x, t*) at indicated fixed times, while the colour maps in (b, d, f) represent *n*(*x, t*) across a temporal interval. The pink arrows point in the direction of increasing time. Parameter values: *r*_*m*_ = 0.3, *r*_*a*_ = 0.2, *κ* = 100 and *ϕ* = 0.2 (a, b), *ϕ* = 0.5 (c, d) and *ϕ* = 0.85 (e, f).

In Fig. 1(e, f), we present *n* and *v*_*n*_ when *ϕ* = 0.85. After a period of transient growth from the initial data, *n*(*x, t*) exhibits a patterned solution comprising multiple regions of high density (shown in yellow), which we term cell peaks. These peaks are sharply separated by regions of low cell density (shown in blue). The maximal volume fraction of all cell peaks is approximately equal to *n*(*L, t*), where from (9) we have *n*(*L, t*) ~ *n*_∞_ for *t* ≫ 1. As highlighted by the velocity profiles, cells in low density regions migrate up gradients of *n* toward regions of high cell density, due to attractive forces experienced between them when Σ(*n*) *<* 0. This attraction results in the contraction of the tumour, and consequently the eventual extinction of the tumour. Nevertheless, the patterned solutions observed prior to extinction retain a degree of biological relevance. For example, and as described in a similar multiphase model of tissue growth (Green et al., 2009), the inherent structural instability of a suspension of *in vitro* tumour cells with a spatially-patterned structure could lead to its break-up, and consequently the formation of separate tumour spheroids. Parameter regimes in which we expect pattern formation, that could be indicative of break-up, are presented in Section 4 by determining the instability of travelling-wave solutions.

## 4 Travelling-wave solutions and stability analysis

In this section, we use travelling-wave analysis to obtain the speed at which the tumour either advances or retreats in terms of the model parameters. We also present regions of parameter space in which travelling-wave solutions are linearly unstable in *t*, which indicate when patterned solutions are expected.

### 4.1 Formulation

We write the MBM in terms of the coordinates 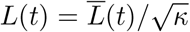 and 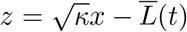, where 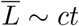 and *z* ∈ (−∞, 0]. Here, *c* is the scaled wave speed, i.e. the speed of travelling waves observed in a simulation of the MBM is 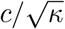. Setting *n* ≡ *n*(*z, t*) and 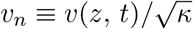, we obtain

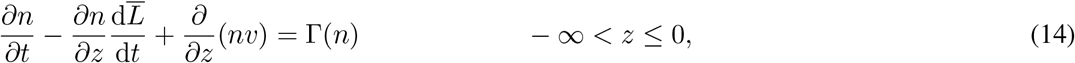

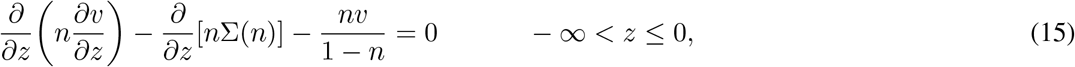

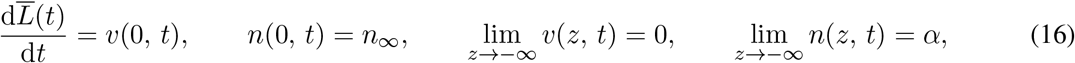

where *α* = (*r*_*m*_ − *r*_*a*_)*/r*_*m*_. We note that since *κ* does not appear in (14)**–**(16), the value of the cell-liquid drag does not determine the stability or direction of travelling waves.

To determine the stability of travelling waves, we introduce the perturbations

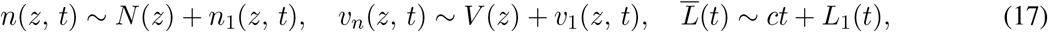

where |*n*_1_|, |*v*_1_|, |*L*_1_| ≪ 1, so that to leading order, (14)**–**(16) provide the system of travelling-wave ODEs

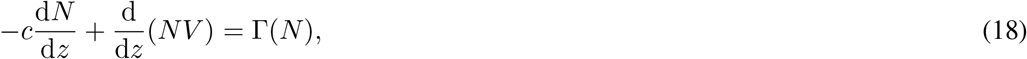

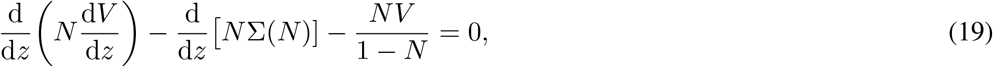

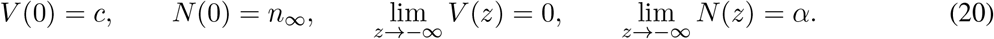

To obtain numerical approximations of *N* (*z*), *V* (*z*) and *c*, we write (18) and (19) as system of three first-order ODEs and use the function `bvp5c` in MATLAB to solve the resulting system. Details of the numerical methods used are presented in Appendix A. In view of the perturbations from (17), the system (14)**–**(16) provides the linearised problem

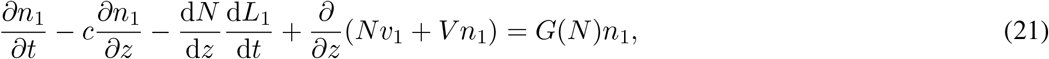

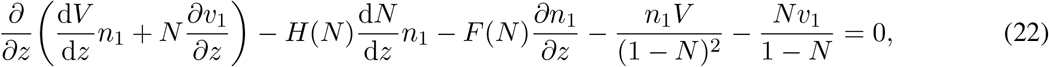

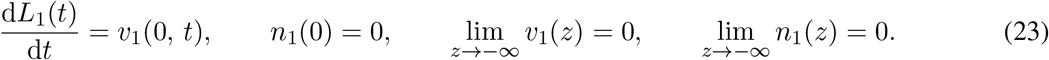

Here, 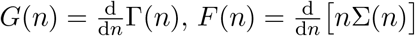 and 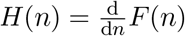, where Γ(*n*) and Σ(*n*) are stated in (1) and(4), respectively.

To determine whether the perturbations *n*_1_, *v*_1_ and *L*_1_ grow or decay with *t*, we set

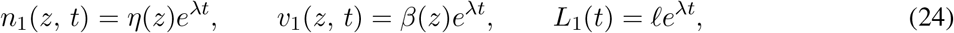

where *ℓ* is a constant and *λ* is the growth rate. If ℜ(*λ*) *>* 0 or ℜ(*λ*) *<* 0, then the solution triple (*N, V, c*) is unstable or stable in *t*, respectively. Substituting (24) into (21)**–**(23), we obtain the autonomous system of ODEs

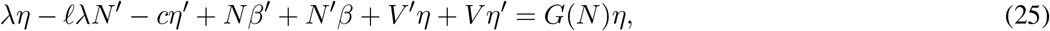

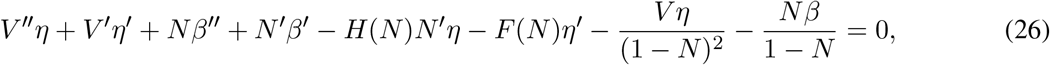

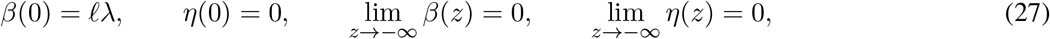

where 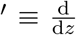. To obtain *λ*, we discretise (25)–(27) using finite differences and assemble the resulting linear system into an eigenvalue problem for *λ*. Using the function `eig` in MATLAB, we obtain the corresponding set of eigenvalues and isolate the pair with the largest real part, Λ = max [ℜ(*λ*)]. If Λ *>* 0, travelling-wave solutions are unstable in *t*, and the MBM may exhibit patterned solutions. If Λ *<* 0, we expect the MBM to exhibit stable travelling-wave solutions.

### 4.2 Results

In this subsection, we present and discuss the criteria for pattern formation obtained from (25)**–**(27), as well as the travelling-wave speed obtained from (18)**–**(20). We first describe the results presented in Fig. 2(a). The solid black line represents the neutral stability curve Λ = 0 obtained numerically from (25)**–**(27) as described above, for a fixed value of *r*_*a*_. Regions in which travelling-wave solutions are unstable (Λ *>* 0) or stable (Λ *<* 0) are indicated. In Fig. 2(a), we also present the scaled wave speed *c* when travelling-wave solutions are stable, obtained numerically from (18)**–**(20), via the methods presented in Appendix A. The dashed black line represents the curve *c* = 0, and regions in which the tumour advances (*c >* 0) or retreats (*c <* 0) are indicated. We recall that when travelling-wave solutions are stable, the wave speed, 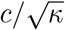, is the speed at which the tumour either advances or retreats.

**Figure 2:**
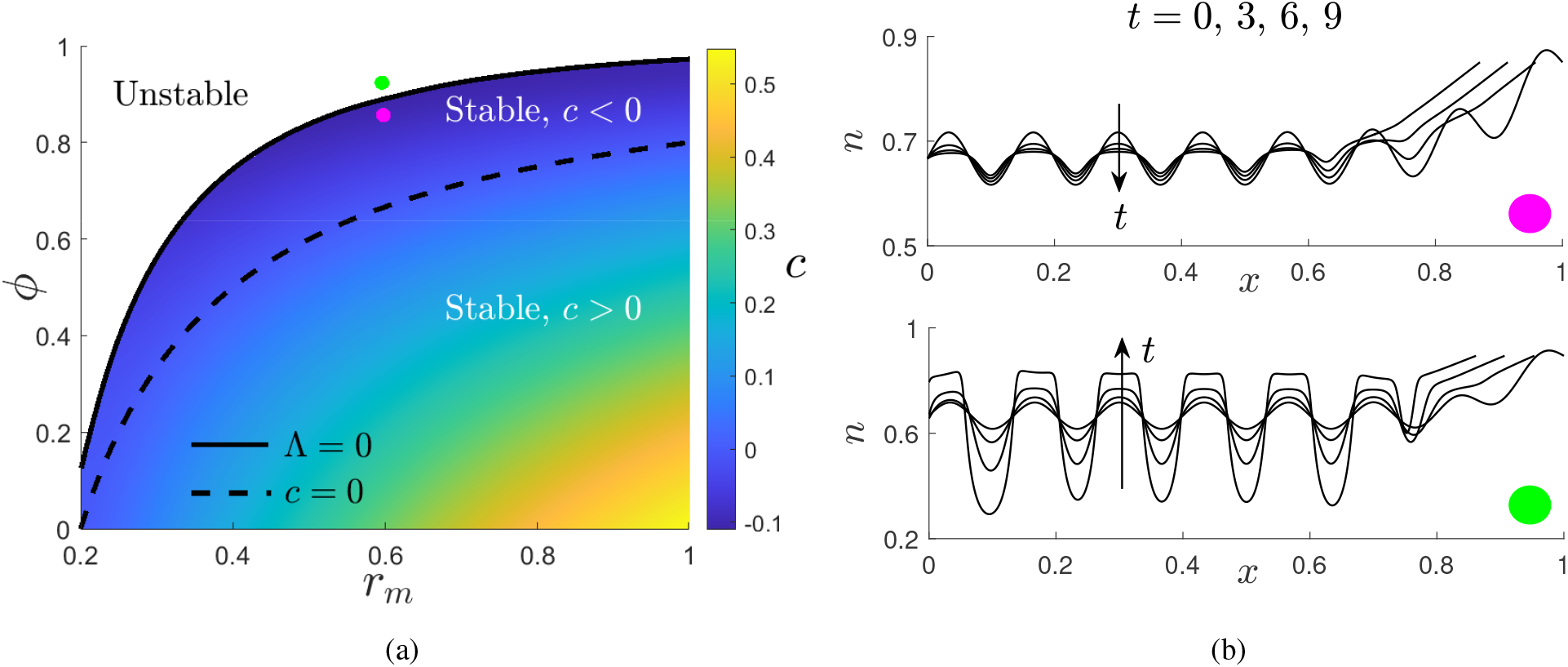
An analysis of travelling-wave solutions and their stability. In (a), the black solid line is the curve Λ = 0 obtained numerically from (25)**–**(27). The black dashed line is the curve *c* = 0. The colour map shows *c*, obtained numerically from (18)**–**(20). Parameter values for (a) are 0 *< ϕ <* 1, 0.2 *< r*_*m*_ *≤* 1 and *r*_*a*_ = 0.2. In (b), we plot *n*(*x, t*) from the MBM for *ϕ* = 0.87 (top panel, pink dot) and *ϕ* = 0.91 (bottom panel, green dot) when *t* = 0, 3, 6, 9. Parameter values for (b) are *r*_*m*_ = 0.6, *r*_*a*_ = 0.2, *κ* = 50. For both panels in (b), we use *n*_*i*_(*x*) = *N* (*x*) + 0.05 sin(15*πx*), where *N* (*x*) is the travelling-wave solution when *t* = 0.

To illustrate the accuracy of the stability regions shown in Fig. 2(a), we compare two numerical solutions of *n*(*x, t*) from the MBM that are marginally stable or unstable. In Fig. 2(a), the green and pink dots indicate two parameter regimes in which travelling-wave solutions are marginally unstable and stable, respectively. In Fig. 2(b), we present *n*(*x, t*) from the MBM corresponding to these two parameter regimes at exemplar early timepoints. For each parameter regime, we take the initial conditions to be a perturbation of the travelling-wave solution *N* obtained from (18)**–**(20), i.e.

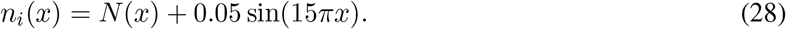

When the marginally stable parameter regime (pink dot) is used, solutions decay from the initial perturbation toward a backward-moving travelling wave, similar to that shown in Fig. 1(c). As expected, solutions grow toward a pattern-forming solution from the initial perturbation when the marginally unstable parameter regime (green dot) is used. Hence, the results presented in Fig. 2(b) are consistent with the unstable and stable regions shown in Fig. 2(a), and illustrate a good level of accuracy of the neutral stability curve, Λ = 0.

As seen from the neutral stability curve Λ = 0 in Fig. 2(a), the value of *ϕ* required to destabilise travelling-wave solutions decreases with *r*_*m*_. This suggests that patterns will form if there is insufficient cell growth to support the uniform migration of cells towards their natural packing density. Numerical solutions of the travelling-wave ODEs from (18)**–**(20) indicate that the sign of *c* corresponds to the sign of *α* − *ϕ*. This is consistent with the results in Fig. 2(b), as well as the slow-wave (|*c*| ≪ 1) asymptotic analysis of (18)**–**(20) given in Appendix A.2. When *c >* 0 (corresponding to when *α > ϕ*), we have *n > ϕ* behind the wave-front and cells are in a constant state of repulsion. Consequently, the tumour edge expands to provide space to which the cells can migrate. Conversely, when *c <* 0 (corresponding to when *α < ϕ*), we have *n < ϕ* behind the wave-front and cells are in a constant state of attraction which results in the contraction of the tumour edge.

Fig. 2(a) indicates that *c* is greatest when *r*_*m*_ = 1 and *ϕ* → 0. This suggests that the tumour grows quickest when the rate of cell production is large, but the cells’ natural packing density is small (thereby increasing the strength of repulsive forces experienced between cells when *n > ϕ*).

## 5 The stability of a spatially-uniform steady state

In contrast to the travelling-wave stability analysis presented in the previous section, we now determine the stability of the spatially-uniform steady state satisfying equations (1) and (3) on an infinite spatial domain. This stability analysis neglects the effects of the boundary conditions at the tumour edge, and has been used informally in similar moving boundary models of tissue growth, e.g Lemon and King (2007) and Byrne et al. (2003). A comparison between this stability analysis and that from Section 4, where the effects of the moving boundary are incorporated, allows us to identify whether the moving boundary at the tumour edge affects the onset of pattern formation.

The non-trivial, spatially-uniform steady state of (1) and (3) is given by

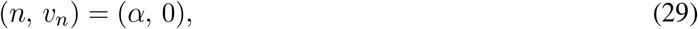

where *α* = (*r*_*m*_ − *r*_*a*_)*/r*_*m*_. We introduce the linearisation *n* ~ *α* + *n*_*p*_(*x, t*) and *v*_*n*_ ~ *v*_*p*_(*x, t*), where |*n*_*p*_|, |*v*_*p*_| ≪ 1 satisfy the linearised problem

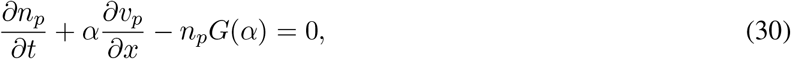

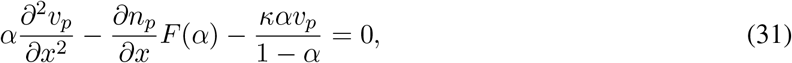

where 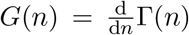 and 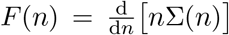. To determine whether the perturbations *n*_*p*_ and *v*_*p*_ grow or decay in *t*, we set (*n*_*p*_, *v*_*p*_) ∝ *e*^*iγx*+*ρt*^ where *γ* is the perturbation wave number and *ρ* is the growth rate. Substituting this ansatz into (30) and (31), we obtain the dispersion relation

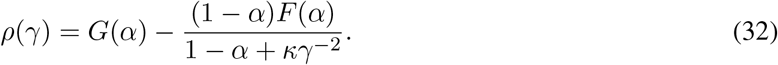

If *ρ >* 0 or *ρ <* 0, then the steady state from (29) is unstable or stable, respectively. As such, it is useful to consider the quantity

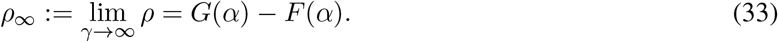

If *ρ*_∞_ *>* 0, then *ρ*(*γ*) is monotonically increasing and since *ρ*(0) *<* 0, we have *ρ*(*γ*) *>* 0 for a sufficiently large value of *γ*. If *ρ*_∞_ *<* 0, then *ρ*(*γ*) is monotonically decreasing so that *ρ*(*γ*) *<* 0. Therefore, the steady state from is unstable or stable when *ρ*_∞_ *>* 0 or *ρ*_∞_ *<* 0, respectively. In Fig. 3(a), we illustrate the function *ρ*(*γ*) for two different parameter regimes which are provided in the caption, which give rise to the cases *ρ*_∞_ *>* 0 and *ρ*_∞_ *<* 0.

**Figure 3:**
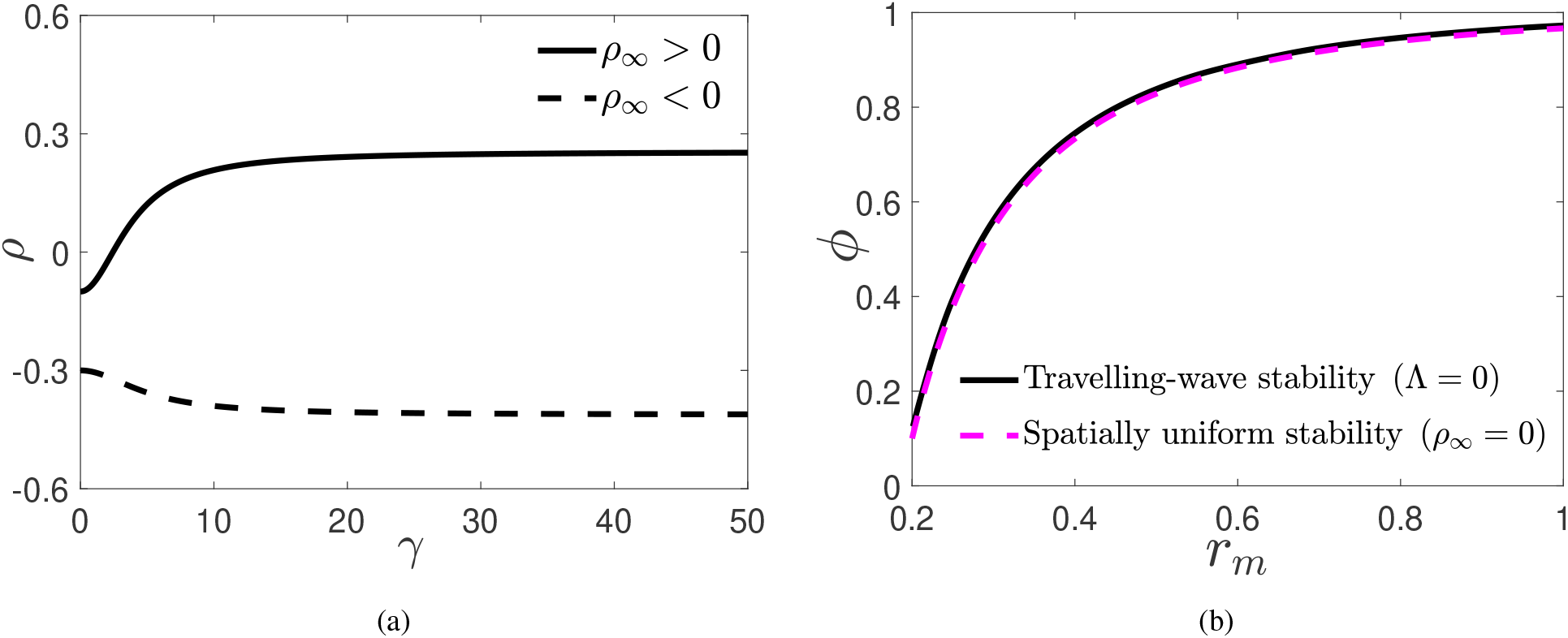
A stability analysis of the spatially-uniform steady state (*n, v*_*n*_) = (*α*, 0). In (a), the solid black line is the function *ρ*(*γ*) from (32) for *r*_*m*_ = 0.3, *r*_*a*_ = 0.2 and *ϕ* = 0.7, whilst the dashed black line is *ρ*(*γ*) for *r*_*m*_ = 0.5, *r*_*a*_ = 0.2 and *ϕ* = 0.7. In (b), the solid black line is the curve Λ = 0 obtained numerically from (25)**–**(27), whilst the dashed pink line in is the curve *ρ*_*∞*_ = 0, both with *r*_*a*_ = 0.2.

We now compare the stability analysis presented in this section with that from Section 4. In Fig. 3(b), we compare the neutral travelling-wave stability curve Λ = 0 obtained numerically from (25)**–**(27) with the neutral stability curve of the spatially-uniform steady-state *ρ*_∞_ = 0 obtained in this section. As seen in Fig. 3(b), these two approaches to computing the respective neutral stability curves are in good agreement. This suggests that the values of cell production, cell death and the natural packing density largely determine the emergence of patterned solutions, in contrast to the inclusion or exclusion of the moving boundary. We therefore expect the destabilising mechanism giving rise to patterned solutions to be the attractive forces experienced by cells occurring when *n < ϕ*. The good agreement between the curves presented in Fig. 3(b) justifies using the simpler analysis described in this section to determine criteria for spatial patterning, rather than the more computationally exhaustive analysis described in Section 4.

The close agreement between the neutral stability curves presented in Fig. 3(b) allows us to exploit the relation from (29) to deduce how the model parameters affect the onset of pattern formation. The parameter regime in which (29) is unstable, and hence patterned solutions are expected, can be found explicitly from *p*_∞_ > 0, i.e.

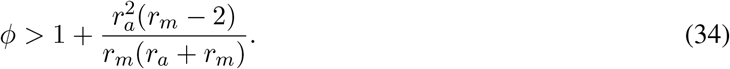

Since 0 *< ϕ <* 1, (34) indicates that solutions will not exhibit pattern formation when *r*_*m*_ ≥ 2. This indicates that a high cell proliferation rate alone can curtail the formation of patterned solutions, regardless of the strength of attractive forces experienced between cells. Furthermore, (34) suggests that no patterns will form if there is a negligible rate of cell death. We also note that (34) is independent of *κ*, this being consistent with the analysis from Section 4, which indicated that the value of the cell-liquid drag does not determine whether solutions will exhibit pattern formation.

## 6 Qualitative analysis of pattern formation

In this section, we examine the qualitative features of patterned solutions like those shown in Fig. 1(f), which are formed by multiple regions of high cell density (termed cell peaks), separated by regions of low cell density. As discussed in Section 3, the inherent structural instability of a suspension of *in vitro* tumour cells with a patterned structure could lead to its break-up, and consequently the formation of separate tumour spheroids. Whilst tumour break-up is not explicitly modelled here, a qualitative analysis of patterned solutions nonetheless provides insight into how the value of model parameters could affect the onset of such a process.

### 6.1 Mathematical formulation

In this subsection, we define the metrics used to quantify the number and width of cell peaks. We define the number of cell peaks *β*(*t*), to be the number of intervals within *x* ∈ [0, *L*(*t*)] such that *n*(*x, t*) is greater than a threshold V. The set containing these intervals, *X*(*t*), is defined as

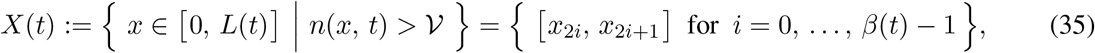

so that *β*(*t*) is the number of intervals at time *t*. Fig 4(a) shows an example patterned solution illustrating these intervals for a fixed value of *t* and 𝒱, where we observe four cell peaks. We define the width of a cell peak occupying [*x*_2*i*_, *x*_2*i*+1_] to be *x*_2*i*+1_ − *x*_2*i*_. Since there are often multiple cell peaks, it is informative to obtain the average width of all cell peaks at any *t*. We define this average to be

**Figure 4:**
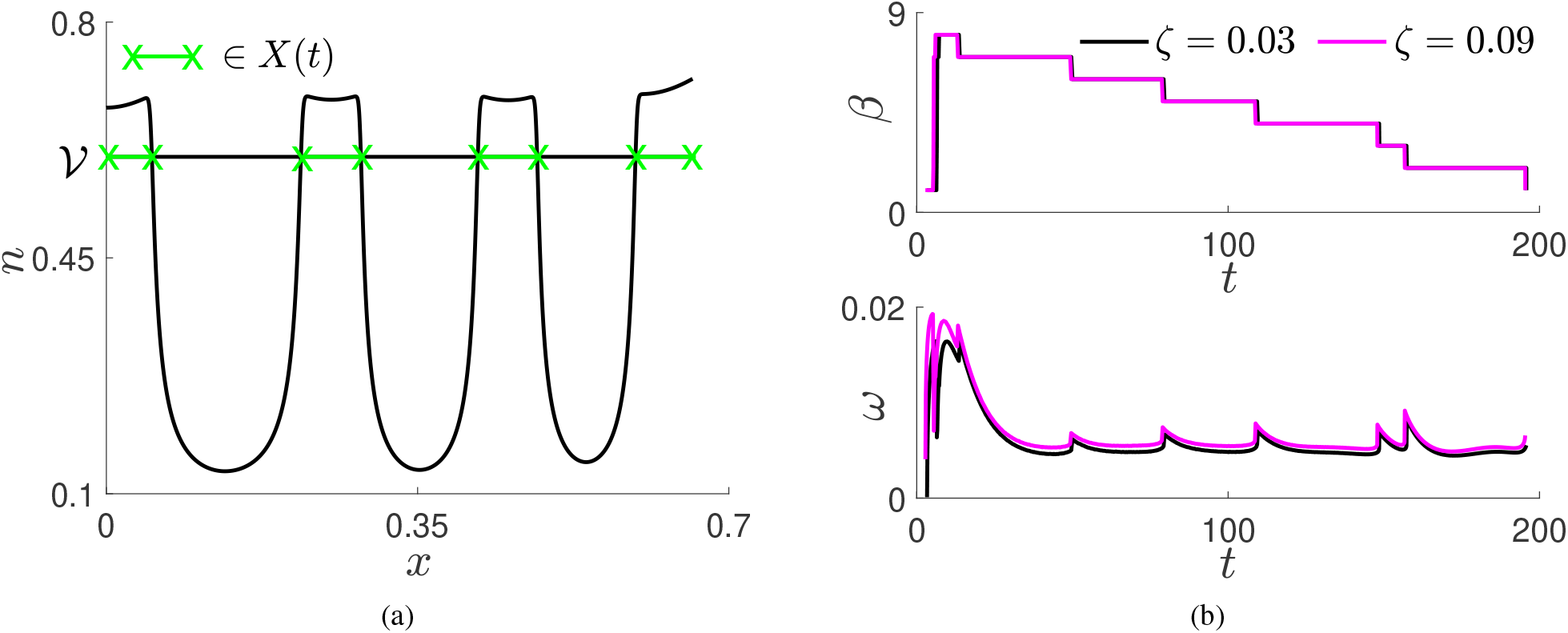
A quantification of patterned solution features. In (a), we present a numerical solution *n*(*x, t*) from the MBM, equations (1), (3), (5)**–**(8), at *t* = 40 for *r*_*m*_ = 0.4, *r*_*a*_ = 0.2, *ϕ* = 0.75, *κ* = 25 and *n*_*i*_(*x*) = *α* + 0.05 sin(15*πx*). The green line segments between the green crosses indicate the elements in *X*(*t*) from (35). In (b), we present the metrics *β*(*t*) and *ω*(*t*) for *ζ* = 0.03 and *ζ* = 0.09. We use *r*_*m*_ = 0.3, *r*_*a*_ = 0.2, *ϕ* = 0.85, *κ* = 100 and *n*_*i*_(*x*) defined as in (a)

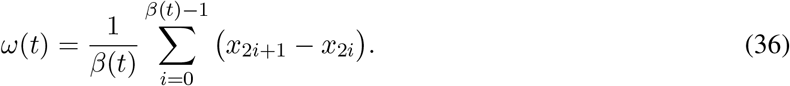

Furthermore, we restrict the *t*-dependent metrics *β*(*t*) and *ω*(*t*) to *t* ∈ [*T*_1_, *T*_2_], where *T*_1_ and *T*_2_ are the times at which the first and last single cell peak are observed, respectively.

Naturally, the choice of the threshold V can influence the metrics *β*(*t*) and *ω*(*t*). As discussed in Section 3, the maximal volume fraction of cell peaks, such as those shown in yellow in Fig. 1(f), is approximately (but never exceeds) the value of *n*_∞_. To capture the features of fully formed cell peaks, we therefore set V = *n*_∞_−*ζ*, where *ζ >* 0 is small. In Fig. 4(b), we present *β*(*t*) and *ω*(*t*) on *t* ∈ [*T*_1_, *T*_2_], computed from the numerical solution of *n*(*x, t*) presented in Fig. 1(f), for two different values of *ζ*. The negligible difference between the metrics for the two values of *ζ* suggests that *β*(*t*) and *ω*(*t*) are not sensitive to the value of *ζ* selected.

In order to examine how the model parameters affect the qualitative features of patterned solutions, it is instructive to associate the metrics *β*(*t*) and *ω*(*t*) with scalar quantities. We define

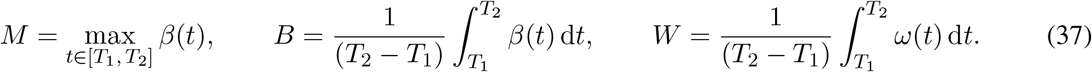

Here, *M* describes the maximum number of cell peaks, whilst the time-averages *B* and *W* describe the average number and average width of cell peaks, in a given simulation of the MBM for *t* ∈ [*T*_1_, *T*_2_]. Additionally, we take *n*_*i*_(*x*) = *α* + 0.05 sin(*νπx*) where *ν* is a positive integer. By varying *ν*, we can investigate how the initial cell distribution affects the qualitative features of patterned solutions.

### 6.2 Results

We now use the quantities described in (37) to investigate how the model parameters affect the qualitative features of patterned solutions. In Figs. 5, 6 and 7, we present the three quantities (*B, M, W*) as functions of *κ, ϕ* and *ν*, respectively. In each of these figures, we also include numerical solutions of *n*(*x, t*) from the MBM for three exemplar values of the respective parameter, as indicated by the coloured dots.

**Figure 5:**
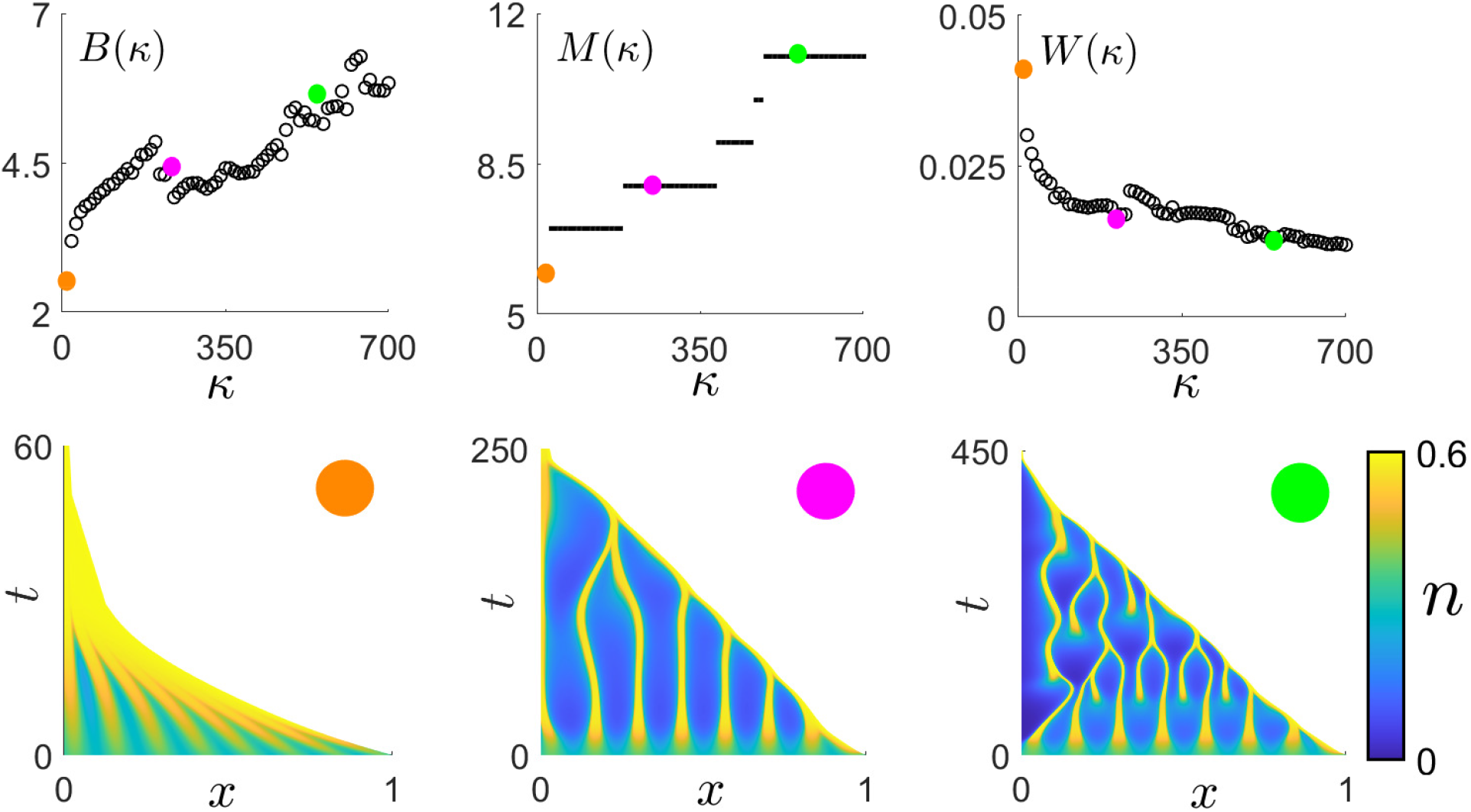
The variation of *B, M* and *W* from (37) with *φ*, computed from numerical solutions of the MBM, together with exemplar numerical solutions of *n*(*x, t*) at *κ* = 5 (orange dot), *κ* = 200 (pink dot) and *κ* = 500 (green dot). Remaining parameter values: *r*_*m*_ = 0.3, *r*_*a*_ = 0.2, *ϕ* = 0.65, *ν* = 15.

**Figure 6:**
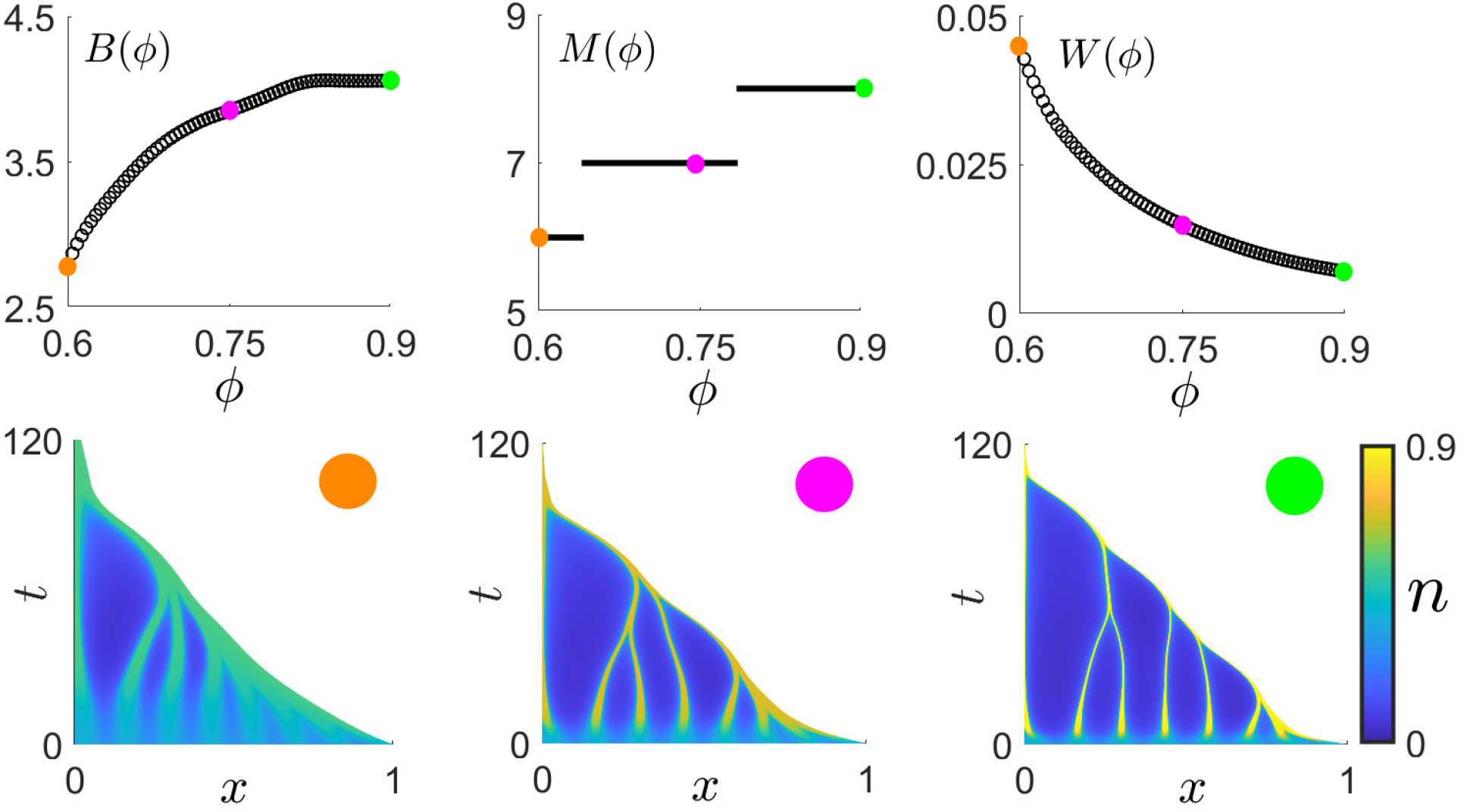
The variation of *B, M* and *W* from (37) with *ϕ*, computed from numerical solutions of the MBM, together with exemplar numerical solutions of *n*(*x, t*) at *ϕ* = 0.6 (orange dot), *ϕ* = 0.75 (pink dot) and *ϕ* = 0.9 (green dot). Remaining parameter values: *r*_*m*_ = 0.3, *r*_*a*_ = 0.2, *κ* = 25, *ν* = 15.

**Figure 7:**
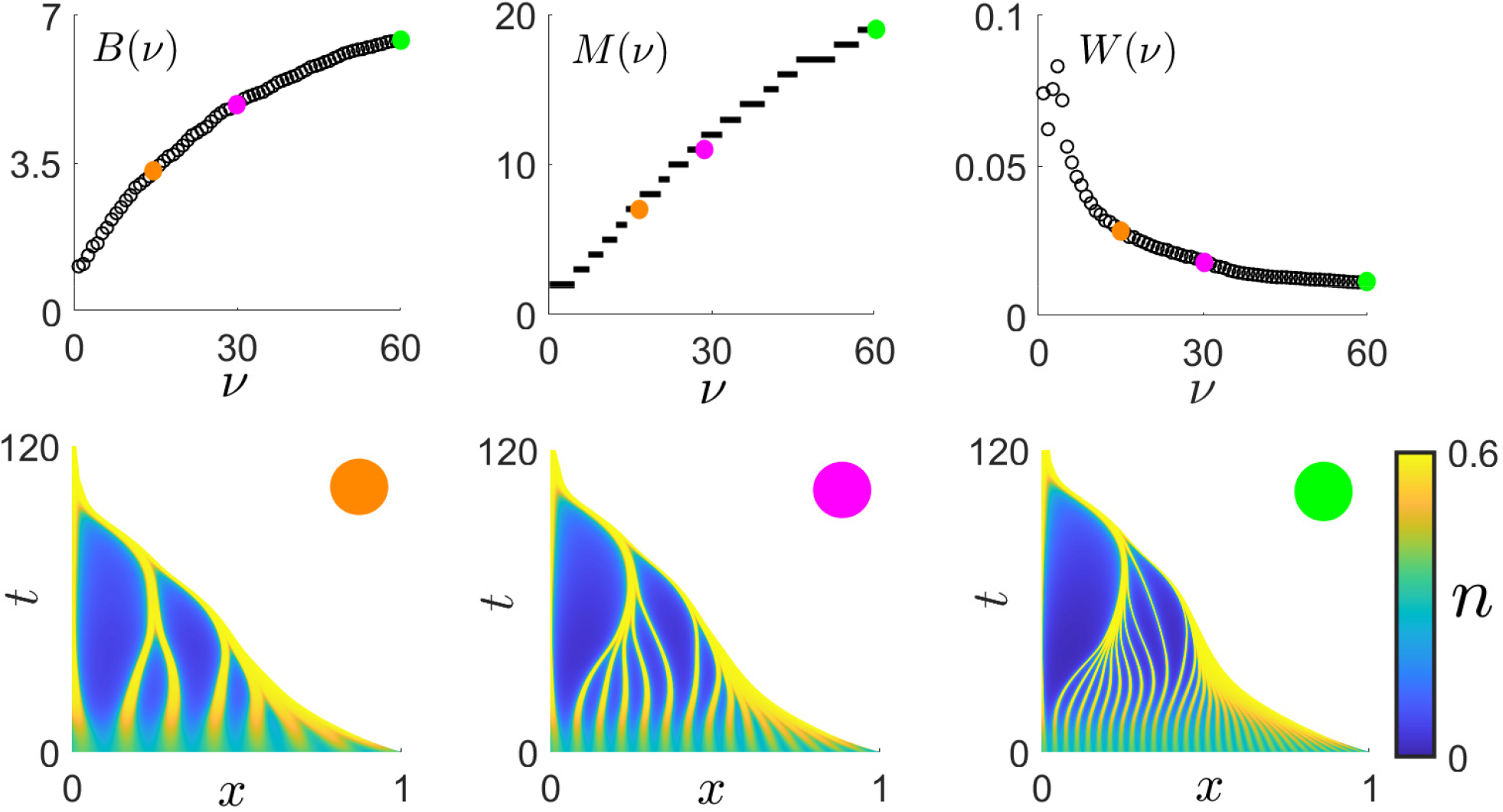
The variation of *B, M* and *W* from (37) with *ν*, computed from numerical solutions of the MBM, together with exemplar numerical solutions of *n*(*x, t*) at *ν* = 15 (orange dot), *ν* = 30 (pink dot) and *ν* = 60 (green dot). Remaining parameter values: *r*_*m*_ = 0.3, *r*_*a*_ = 0.2, *ϕ* = 0.65, *κ* = 25.

The numerical solutions of *n*(*x, t*) presented in Figs. 5 and 7 suggest that the values of *κ* and *ν* have a significant effect on the number of cell peaks, and this observation is substantiated by the observed range over which the quantities *B*(*κ*) & *M* (*κ*) and *B*(*ν*) & *M* (*ν*) increase. Furthermore, the numerical solutions of *n*(*x, t*) and the quantities *B*(*ϕ*) and *M* (*ϕ*) presented in Fig. 6 indicate that *ϕ* does not greatly influence the number of cell peaks. Interestingly, this observation indicates that the strength of attractive forces experienced by cells when *n < ϕ* is not indicative of the number of separate spheroids that result from the break-up of a suspension of *in vitro* tumour cells.

As highlighted by the metric *W* (*ϕ*) and the numerical solutions of *n*(*x, t*) presented in Fig. 6, the width of the cell peaks decreases as *ϕ* increases, as a result of the available cells migrating toward a larger natural packing density. In contrast to this, the quantities *W* (*κ*) and *W* (*ν*) presented in Figs. 5 and 7 decrease due to the available cells being distributed over a larger number of cell peaks. In particular, the value of *ν* has the most significant effect on the cell peak width when compared with the effects of varying *κ* and *ϕ*. Whilst the number of cell peaks increase with *κ* and *ν* (as discussed above), the resulting decrease of cell peak width may have implications on the viability of new tumour spheroids that result from *in vitro* break-up.

Whilst the quantities *B*(*κ*) and *B*(*ν*) provide insight into the average number of cell peaks, they do not capture the differences in the qualitative features of patterns seen in Figs. 5 and 7 at different points in time. For example, the patterned solutions in Fig. 7 illustrate that the number of cell peaks observed at early times are determined by the value of *ν*; however, these peaks are not sustained and merge together at a later time. Additionally, the patterned solutions in Fig. 5 indicate that the value of *κ* controls the number of peaks at later times. In particular, the pink-dot panel in Fig. 5 indicates that there exists a relationship between the values of *κ* and *ν* such that the cell peaks generated by the initial cell distribution are sustained throughout the tumour. However, for a sufficiently larger value of *κ*, spontaneous pattern emergence is observed in the regions of low cell density, as illustrated in the green-dot panel in Fig. 5. In the context of *in vitro* tumour growth, these observations suggests that the initial cell distribution may have a limited effect on the number of cell peaks that can break away from a primary suspension of tumour cells to generate tumour spheroids, if there is insufficient drag between the cells and liquid. This observation is in agreement with the qualitative analysis of the multiphase description of in vitro liver cell aggregation presented in Green et al. (2009).

## 7 Conclusion

In this work, we analyse spatially-patterned and travelling-wave solutions of the two-phase, moving boundary model of tumour growth developed in Byrne et al. (2003). The model consists of two equations governing a cell volume fraction denoted by *n* and its associated velocity, as well as a moving boundary condition for the tumour edge. Mechanisms to represent forces generated by cell-cell interactions are accounted for by considering relevant constitutive assumptions in a similar fashion to those in Byrne et al. (2003) and Breward et al. (2002). One important parameter related to cell-cell interactions is *ϕ*, which represents the cells’ natural packing density. If *n > ϕ*, then cells repel each other to relieve membrane stress, and if *n < ϕ*, then cells will attract one another due to their filopodia coming into contact. In keeping with King and Franks (2004), we assume that nutrient is abundantly distributed throughout the tumour. Whilst this assumption omits important elements such as nutrient limited induced cell death, it is physically relevant in the context of an *in vivo* tumour in the initial stage of development where all cells are adequately nourished. Additionally, this nutrient rich assumption is appropriate when considering the initial growth of a suspension of *in vitro* tumour cells (Byrne et al., 2003). Whilst the model developed in Byrne et al. (2003) pertains to both *in vivo* and *in vitro* tumour growth, the techniques developed in its analysis regarding pattern formation can be applied to a wider class of multiphase tissue growth models, such as that described in Lemon and King (2007).

Solutions of the tumour model analysed herein can develop into a forward- or backward-moving travelling wave, which correspond to the growth or retreat of the tumour edge, the latter resulting in tumour extinction. A travelling-wave analysis is used to obtain criteria for the growth or extinction of the tumour via stable travelling waves, in terms of model parameters. We determine that forward-moving travelling-wave solutions have *n > ϕ* behind the wave-front, and cells are in a constant state of repulsion. Consequently, the tumour edge expands to provide space in which the cells can migrate. Conversely, if *n < ϕ*, cells behind the wave-front are in a constant state of attraction, which results in the contraction of the tumour edge and a backward-moving travelling wave. We find that the tumour grows the fastest when the rate of cell production is large, but the cells’ natural packing density is small (thereby increasing the strength of repulsive forces experienced between cells when *n > ϕ*). We also find that the value of the cell-liquid drag does not determine the direction of travelling waves, but does significantly control the wave speed.

We also observe patterned solutions that are associated with multiple regions of high cell density (termed cell peaks) separated by regions of low cell density. From an initial cell distribution, cells migrate up gradients of *n* from regions of low to high density to form distinct cell peaks, due to attractive forces experienced between them when *n < ϕ*. This attraction results in the contraction of the tumour, and consequently the eventual extinction of the tumour. Nevertheless, patterned solutions observed prior to extinction retain a degree of biological relevance. For example, and as described in Green et al. (2009), the inherent structural instability of a suspension of *in vitro* tumour with patterned structures could lead to break-up, and consequently the formation of separate spheroids. Whilst the aspect of tumour break-up is not explicitly modelled here, a qualitative analysis of patterned solutions nonetheless provides insight into how model parameters could affect the onset of such a process. Notably, we find that the initial cell distribution only determines the number of cell peaks within the tumour at early times, whereas the value of the cell-liquid drag determines the number of cell peaks at later times. This suggests that the initial cell distribution could have a limited effect on the number of new spheroids that result from *in vitro* tumour break-up, which is in agreement with the qualitative analysis of the multiphase description of *in vitro* liver cell aggregation presented in Green et al. (2009).

We determine the instability of travelling-wave solutions in time to obtain regions of parameter space in which patterned solutions are observed. This stability analysis incorporates the effects of the boundary conditions imposed at the tumour edge and tumour core, as well as the moving tumour edge. The accuracy of the stability regions generated by this linear stability analysis are verified by comparing them with numerical solutions of the cell volume fraction in suitable parameter regimes. The biological implications of the travelling-wave stability analysis suggest that pattern solutions will emerge if there is insufficient net cell growth to support the attraction of cells uniformly toward their natural packing density.

In addition to the travelling-wave stability analysis described above, we determine the stability of a spatially-uniform steady state. In contrast to the travelling-wave stability analysis, this spatially-uniform steady state does not satisfy the moving boundary condition at the tumour edge. Interestingly, however, the regions of instability obtained via the travelling-wave and spatially-uniform stability analyses are in very good agreement, suggesting that the inclusion or exclusion of the moving boundary does not determine when patterned solutions will form. We therefore expect the destabilising mechanism giving rise to pattern formation to be the attractive forces experienced by cells occurring when *n < ϕ*. A similar multiphase moving boundary model to that described in this paper is presented in Lemon and King (2007), and the stability of a spatially-uniform steady state (which does not satisfy the moving boundary condition) is analysed to obtain criteria in which pattern-forming solutions are expected. The good agreement between the two types of stability analyses presented in this paper provides a justification for exploiting the simpler spatially-uniform analysis, such as for the class of multiphase models in Lemon and King (2007).

A possible extension of the work presented in this paper is to investigate how additional constitutive assumptions describing cell-cell interactions affect the onset of pattern formation. For example, in Breward et al. (2002), a mechanism is employed in the constitutive assumptions that ensures attractive forces experienced between cells are short-range, so that cells do not attempt to cluster if they are too far apart. Another natural extension of this work is to examine the influence that nutrient limitation has on the onset of pattern formation within a tumour. In contrast to the model presented in this paper, preliminary numerical simulations of the nutrient-limited extension indicate that pattern-forming solutions can exist on a growing spatial domain.

## Acknowledgements

Jacob M. Jepson acknowledges funding provided by the Engineering and Physical Sciences Research Council (Grant Number EP/R513283/1). We are grateful to Dr Andrew Krause and Professor John King for helpful discussions.

## A Numerical methods for travelling-wave solutions

In this section, we present the numerical methods used to determine *N* (*z*), *V* (*z*) and *c* from (18)**–**(20).

### A.1 Formulation of travelling-wave ODEs

For numerical purposes, we express (18)**–**(20) as a system of three, first order ODEs by introducing the variables *U* (*z*) = *V* ^′^(*z*) where ^′^ ≡ d*/*d*z*. We obtain

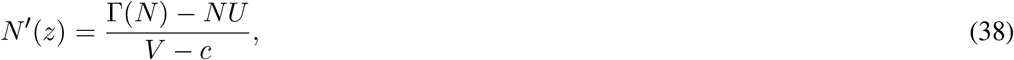

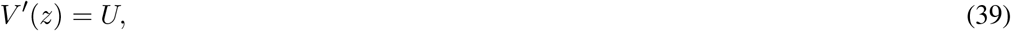

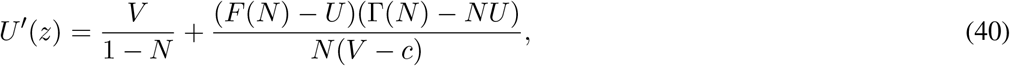

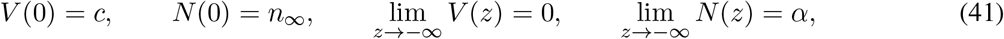

where *α* = 1−*r*_*a*_*/r*_*m*_. The system from (38)**–**(41) is computationally singular at *z* = 0 because of the boundary condition *V* (0) = *c* and the right-hand side of (38) and (40). In view of this, we consider the leading edge of the wavefront to be *z* = −*δ* where *δ* is sufficiently small such that the solution converges. We numerically integrate (38)**–**(41) on *z* ∈ [−*Z*, −*δ*] in MATLAB using the function `bvp5c` which uses a fifth-order, implicit Runge-Kutta method. We impose the boundary conditions from (41) as *z* → −∞ at *z* = −*Z*, where *Z* is sufficiently large such that the solution converges. The Taylor expansions of *N* (*z*) and *V* (*z*) around *z* = 0, *N* (*z*) ~ *n*_∞_ + *N* ^′^(0)*z* and *V* (*z*) ~ *c* + Σ(*n*_∞_)*z*, are used as boundary conditions at *z* = −*δ*. The value of *N* ^′^(0) is found by combining (38) and (41) at *z* = 0 and employing L’Hôpital’s rule. We use the built-in parameter estimation feature in `bvp5c` to obtain the value of *c*.

### A.2 Slow wave speed (|*c*| ≪ 1) asymptotic analysis

The function `bvp5c` requires an initial approximation of the solutions *N, V* and *c* satisfying (38)**–**(41) in order to converge. For this, we use asymptotic solutions valid when |*α* − *n*_∞_| ≪ 1. When |*α* − *n*_∞_| ≪ 1, then |*c*| ≪ 1, and we linearise *N* and *V* around their far-field values, 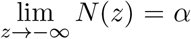 and 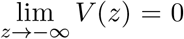. Following Lemon and King (2007), we introduce the perturbations *N* ~*α*+*εN*_1_, *V* ~ *εV*_0_ and *c*~ *εc*_0_ where *ε* = *α* − *n*_∞_ and |*ε*| ≪ 1. The perturbations *N*_1_ and *V*_0_ satisfy

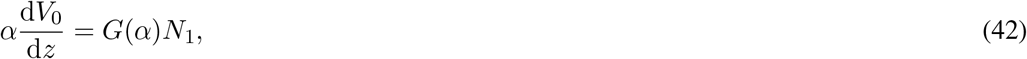

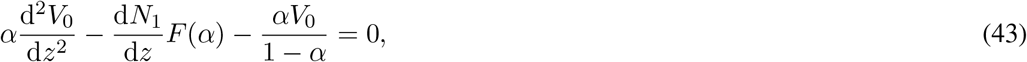

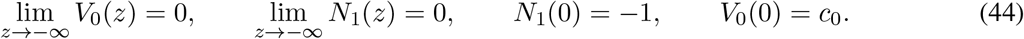

where 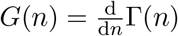 and 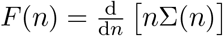. Combining (42) and (43) provides

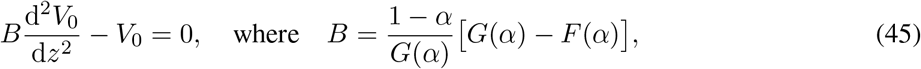

so that 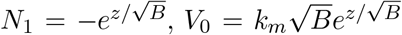 and 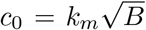. These asymptotic solutions are used as initial guesses of *N, V* and *c* from (38)**–**(41). For more general parameter values, such as those presented in Fig. 2(a), the method of parameter continuation is used.

## References

A. Boonderick, W. Triampo, and N. Nuttavut. A Review of Cellular Automata Models of Tumor Growth. International Mathematics forum, 2010. 61:3023–3029.

C. J. W. Breward, H. M. Byrne, and C. E. Lewis. The role of cell-cell interactions in a two-phase model for avascular tumour growth. Journal of Mathematical Biology, 2002. 45:125–152.

A. C. Burton. Rate of growth of solid tumours as a problem of diffusion. Growth, 1966. 30(2):157–76.

H. M. Byrne, J.R. King, D.L.S McElwain, and L. Preziosi. Two-Phase Model of Solid Tumour Growth. Applied Mathematics Letters, 2003. 16:567–573.

E. J. Crampin, W. W. Hackborn, and P. K. Maini. Pattern Formation in Reaction–Diffusion Models with Nonuniform Domain Growth. Bulletin of Mathematical Biology, 2002. 64:747–769.

N. P. Fam, S. Verma, M. Kutryk, and D. J. Stewart. Clinician guide to angiogenisis. Circulation, 2003. 108:2613–2618.

S. J. Franks and J. R. King. Interactions between a uniformly proliferating tumour and its surroundings: uniform material properties. Mathematical Medicine and Biology, 2003. 20: 47–89.

R. A. Van Gorder, V. Klika, and A. L. Krause. Turing conditions for pattern forming systems on evolving manifolds. Journal of Mathematical Biology, 2021. 82:4.

J. E. F. Green, S. L. Waters, K. M. Shakesheff, and H. M. Byrne. A Mathematical Model of Liver Cell Aggregation In Vitro. Bulletin of Mathematical Biology, 2009. 71:906–930.

J. E. F. Green, J. P. Whiteley, J. M. Oliver, H. M. Byrne, and S. L. Waters. Pattern formation in multiphase models of chemotactic cell aggregation. Journal of Mathematical Biology, 2018. 35(3):319–346.

H. P. Greenspan. Models for the Growth of a Solid Tumor by Diffusion. Studies in Applied Mathematics, 1972. 52:317–340.

A. V. Hill. The diffusion of oxygen and lactic acid through tissues. Proceedings of the Royal Society B, 1928. 104:39–96.

J. M. Jepson, Reuben D. O’Dea, and Nabil. T Fadai. Travelling-wave and asymptotic analysis of a multiphase moving boundary model for engineered tissue growth. Bulletin of Mathematical Biology, 2022. 84:87.

J. R. King and S. J. Franks. Mathematical analysis of some multi-dimensional tissue-growth models. European Journal of Applied Mathematics, 2004. 15: 273–295.

A. L. Krause, M. A. Ellis, and R. A. Van Gorder. Influence of curvature, growth, and anisotropy on the evolution of Turing patterns on growing manifolds. Bulletin of Mathematical Biology, 2018. pp. 1–41.

G. Lemon and J. King. Travelling-wave behaviour in a multiphase model of a population of cells in an artificial scaffold. Journal of Mathematical Biology, 2007. 24(1):57–83.

C. K. Macnamara. Biomechanical modelling of cancer: Agent-based force-based models of solid tumours within the context of the tumour microenvironment. Computational and Systems Oncology, 2021. DOI: https://doi.org/10.1002/cso2.1018.

D. L. S McElwain and G. J. Pettet. Cell migration in multicell spheroids: Swimming against the tide. Bulletin of Mathematical Biology, 1993. 55:655–674.

J. D. Murray. Mathematical Biology I: An Introduction. Springer-Verlag, 2002.

R. D. O’Dea, S. L. Waters, and H. M. Byrne. A multiphase model for tissue construct growth in a perfusion bioreactor. Mathematical Medicine and Biology, 2010. 27:95127.

A. Patel. Benign vs Malignant Tumors. The Journal of the American Medical Association Oncology, 2020. 6(9):1488.

G. J. Pettet, C. P. Please, M. J. Tindall, and D. L. S McElwain. The Migration of Cells in Multicell Tumor Spheroids. Bulletin of Mathematical Biology, 2001. 63:231–257.

J. Poleszczuk and H. Enderling. A High-Performance Cellular Automaton Model of Tumor Growth with Dynamically Growing Domains. Applied Mathematics (Irvine), 2014. 5(1): 144–152.

G. Sciumè, S. Shelton, W. G. Gray, C. T Miller, F. Hussain, M. Ferrari, P. Decuzzi, and B. A. Schrefler. A multiphase model for three-dimensional tumor growth. New journal of physics, 2013. 15:015005.

R. H. Thomlinson and L. H. Gray. The histological structure of some human lung cancers and the possible implications for radiotherapy. British journal of cancer, 1955. 9:539–549.

K. E. Thompson and H. M. Byrne. Modelling the internalization of labelled cells in tumour spheroids. Studies in Applied Mathematics, 1999. 61:601–623.

G. Toole and M. K. Hurdal. Turing models of cortical folding on exponentially and logistically growing domains. Journal of Dynamics and Differential Equations, 2014. 26, pp. 315–332.

A. Tosin and L. Preziosi. Multiphase modeling of tumor growth with matrix remodeling and fibrosis. Mathematical and Computer Modelling, 2010. 52(7):969–976.

A. Turing. The Chemical Basis of Morphogensis. Philosophical Transactions of the Royal Society of London B, 1952. 237 (641): 37–72.

J. P. Ward and J. R. King. Mathematical modelling of avascular tumour growth. IMA Journal of Mathematics Applied in Medicine Biology, 1997. 14:39–69.

B. R. Zetter. Angiogenesis and tumor metastasis. Annual Review of Medicine, 1998. 49:407–24.

